# Methodology for the identification and analysis of species associations in biological communities using statistical distributions

**DOI:** 10.1101/2024.08.02.606312

**Authors:** Dmitriy G. Seleznev, Alexander A. Prokin, Ekaterina M. Kurina

## Abstract

A methodology for identifying pairs of mutually associated and mutually incompatible species pairs based on qualitative ecological data in the ‘species-samples’ format is described. Statistical criteria for assessing the direction and strength of species associations using discrete hypergeometric and binomial distributions are provided. Additionally, there are proposed methods for visualising selected associations using undirected graphs, combining associated species into groups, and analysing them. An open access web application implementing the described methodology is also presented. The methodology is illustrated with data on 37 alien macrozoobenthic species from 255 samples collected in the Volga River reservoirs.

## Introduction

In nature, the mutual association or incompatibility of species depends on their interspecific relationships, which are the basis for the origination and existence of biological systems. Species can form assemblages, including communities, based on interspecific interactions, which may be direct (competition, predation, parasitism, etc.) or indirect, i.e., mediated or transmitted by a third species or environment (habit facilitation, keystone predation, exploitation, competition, etc.) (Odum & Barrett, 1971; Wootton, 1994; Moon et al., 2010; Sotomayor & Lortie, 2015; etc.).

To study such interactions using species presence-absence matrix data, it is necessary to define pairs of species that are mutually associated or incompatible. At present, ecological studies rarely address this problem. This may be due to the lack of obvious mathematical methods for this purpose.

For this task, in addition to the correlation coefficients and similarity/dissimilarity measures traditionally used in ecology, such as the Jaccard and Ochiai indices (Jaccard, 1912; Ochiai, 1957) or variations of the Sörensen measure (Sörensen, 1948; Bray & Curtis, 1957), some specialised matrix indices have been proposed. For instance, there are the checkerboard index CHECKER (Diamond, 1975), the C-score index (Stone & Roberts, 1990), Schluter’s variance test (Schluter, 1984), and the nestedness temperature (Atmar & Patterson, 1993).

In addition to the necessary normalisation, all of these methods are based on the arithmetic calculation of species co-occurrence in the qualitative matrix. Therefore, the researcher does not have comprehensible criteria for identifying mutually associated species, even such unreliable one as the significance level. In the case of an observed co-occurrence of two species, the significance of the index, as well as the decision on the association of species, directly depends on the total number of samples, which matrix indices often do not take into account. Indeed, having obtained a normalised C-score index of 0.96, the researcher has no formal reason to assert that these species are mutually incompatible and that their low co-occurrence is not due to chance. This issue has been proposed to be resolved by creating a randomly generated null model or a theoretical distribution of the matrix index with the observed species occurrence and the total number of samples (Gotelli, 2000). It is crucial to carefully choose a randomisation algorithm because different permutation schemes can yield vastly different results. The decision about the association or incompatibility of a pair of species is made by comparing the observed index and given confidence intervals of the theoretical distribution. This approach is proposed to be used by the most common ecological R package “vegan” (Oksanen et al., 2019), as well as by the specialised package “EcoSimR” (Gotelli et al., 2015).

In practice, the study of species associations using matrix indices and a simulated null model can encounter certain difficulties. An analysis of 255 samples of alien macrozoobenthic species from Volga River reservoirs with the C-score index and a permutational fixed-equiprobable (FE) null model (Kurina & Seleznev, 2019) revealed 82 positively associated and 5 negatively associated pairs of species. However, 27 positively associated pairs had a normalised C-score index greater than 0.4, reaching a value of 0.55 for the pair of species *Katamysis warpachowskyi* (Sars, 1893) (n_1_=16) and *Pterocuma rostrata* (Sars, 1894) (n_2_=25), which co-occurred in 5 samples.

Therefore, interpreting the normalised C-score index is a challenging task. Furthermore, the tendency of permutational null models to overdiagnose, producing the type I error, has been noted (Sanderson, 2004; Gotelli et al., 2010), as illustrated by the example above.

An estimation of the significance level of indices using a randomisation procedure has a probabilistic nature. This approach may lead to problems with the reproducibility of the results obtained, as well as their instability. In a set of samples with several pairs of species having the same individual and joint co-occurrence, only some of them may show a significant association. Moreover, the set of these pairs may change during repeated modelling.

### Identification of associated pairs of species using statistical distributions

A probabilistic model has recently been proposed for the precise calculation of the probability of a certain event: the co-occurrence of two species in *x* out of *N* samples (Veech, 2013). In that model, the classical definition of probability as a quotient of the number of successful outcomes and the total number of outcomes was used. This formula was subsequently reduced to the probability mass function (PMF) of the hypergeometric distribution (Griffith et al., 2016), which describes the probability of obtaining a specific number of successes in draws without replacement from a finite population. When applied to the co-occurrence of two species, the hypergeometric distribution allows calculations of the probability of finding particular species in samples that already contain other species with a total number of samples *N* and given that species are distributed independently within the community. Let *m* denote the occurrence of a rarer species and *n* denote the occurrence of a more common species. The probability that the species will be observed together precisely in *x* samples of a total of *N* samples is determined by the probability mass function of the hypergeometric distribution:

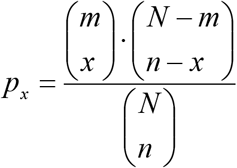

Therefore, the task of determining association or, conversely, incompatibility of two species at a given significance level can be reduced to checking whether the observed joint co-occurrence value falls out of the confidence intervals of the hypergeometric cumulative distribution function (CDF):

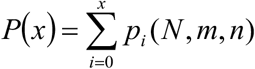

In other words, the hypergeometric CDF allows one to test the null hypothesis that species are independent of each other and that their co-occurrence in samples is random. An alternative is the assumption that the observed positive or negative association of species is not accidental and is due to their biological characteristics or ecological reasons.

It should be noted that the Fisher’s exact test on the contingency table 2×2 (species A, species B; occurrence, co-occurrence) leads to the same result. This is because, as Ronald Fisher points out, such a test for frequency independence leads to a hypergeometric distribution.

A discrete hypergeometric distribution tends to be a binomial distribution with an infinite increase in the general population size (Fig. 1). In this case, a “draws without replacement” model becomes “draws with replacement,” also known as the Bernoulli scheme. When using the hypergeometric distribution, we considered all samples taken as the general population, while samples in which one of the species was found were considered as the draws.

**Figure 1.**
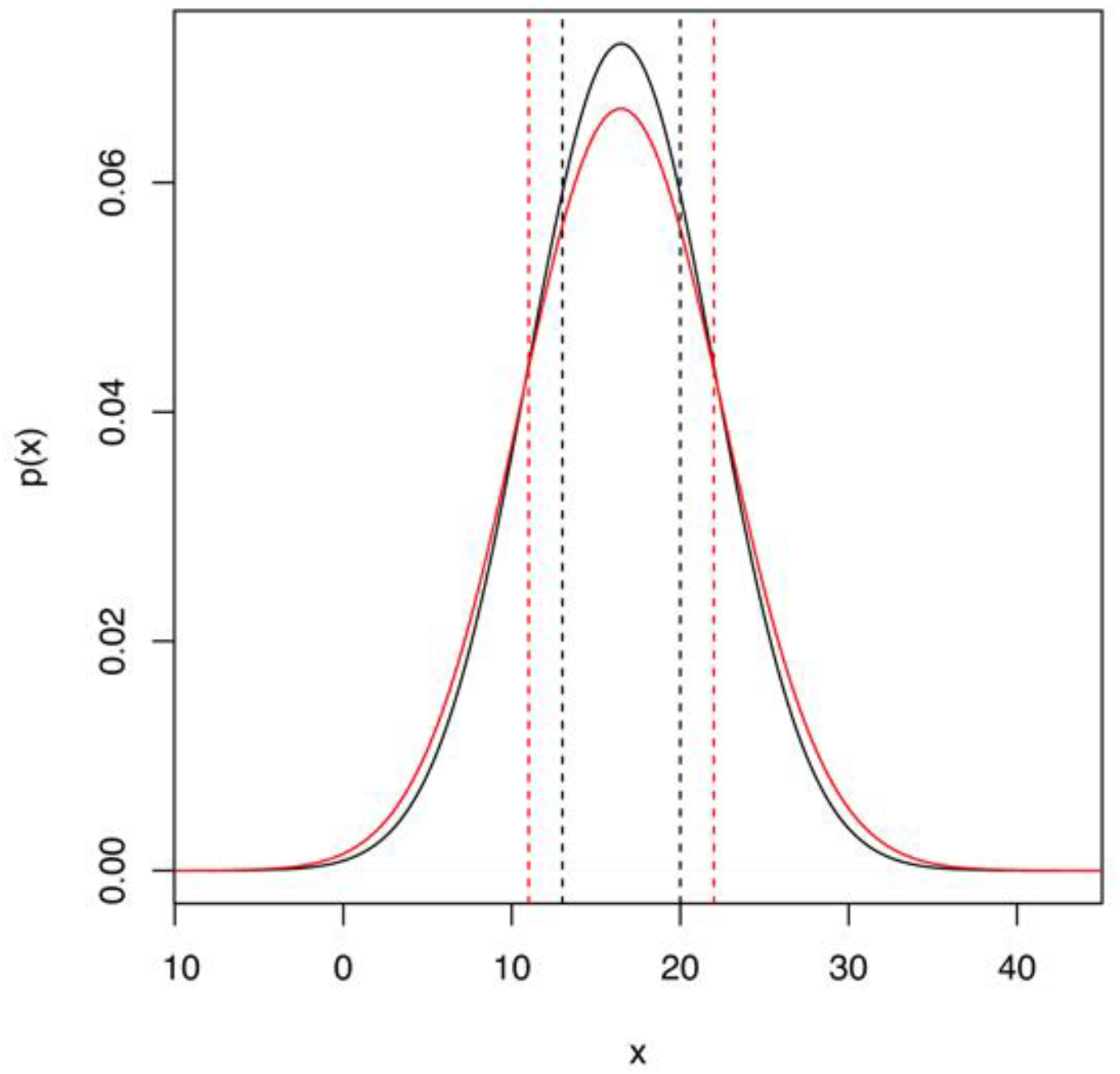
Simulated probability mass function of the hypergeometric (black curve) and binomial (red curve) distributions with 10^6^ simulations of 50 trials and a success probability of 1/3. The vertical lines indicate the boundaries of the 90% confidence intervals.

**Figure 2.**
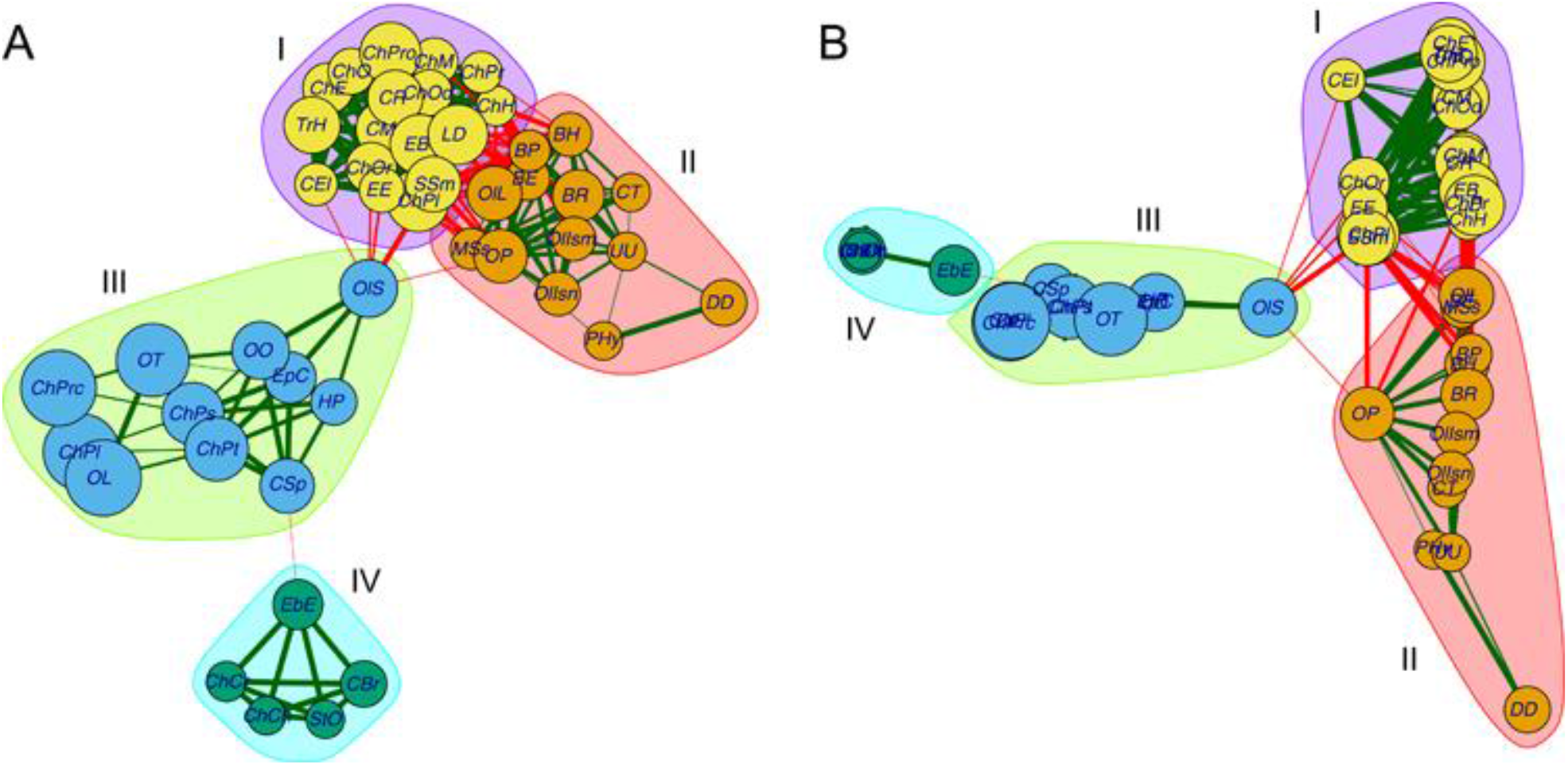
The graph of positively associated pairs of macrozoobenthic species in rivers of the East European Plain. A binomial distribution with a significance level of 0.05 and a multi-level modularity optimisation algorithm for clustering were employed. A – Kamada–Kawai algorithm node layout, B – Multidimensional scaling (MDS) node layout. For species abbreviations, see Golovatyuk & Seleznev (in press).

Let us reformulate the aforementioned conditions. We will consider the entire potentially possible (infinite) number of samples in a given habitat as the general population. The taken samples will be regarded as draws, and the joint co-occurrence of two species in the samples will be defined as draw success. Since we cannot know the a priori probability of success *a priori*, we will use it’s *a posteriori* estimation (Bowers & Brown, 1982; Veech, 2006):

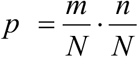

The probability of the joint co-occurrence of two species in *x* samples under these conditions is defined by the probability mass function of the binomial distribution:

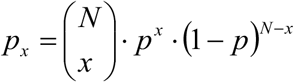

The estimation of the association rate at the given level of significance is also performed by comparing the observed co-occurrence value with the confidence intervals of the binomial cumulative distribution function.

The possibility of making decisions about a probabilistic process based solely on the significance level with a conventional threshold of 0.05 is currently a subject of active debate within the scientific community (see studies published in: The American Statistician, 2019, Vol. 73, Issue Supp 1; Wasserstein & Lazar, 2016). Given that we have defined the distribution function of the probabilistic process a priori, it is possible to determine pairs of mutually associated or incompatible species using an approach based on Bayesian inference rather than the traditional method of testing statistical hypotheses. According to this approach, one or more uniformly distributed characteristics are taken as the evaluated parameter. These may be the occurrence of species or the total number of samples. The probability mass function is then used as a generative function. The posterior distribution, filtered by the observed values of the evaluated parameters, allows us to estimate the probability of the joint co-occurrence of two species. On this basis, we can make one of three decisions: both species occur together by chance, both species are associated, or these two species are incompatible (Fig. 4). It should be noted that the application of Bayesian inference may lead to issues with the reproducibility and stability of the results due to the use of a random variable as a prior distribution, as is the case with the use of randomisation procedures.

The analysis can also use a transposed matrix with initial qualitative data in the “samples-species” format. As a result of the analysis, we obtain a list of pairs of samples whose similarity or difference in species composition significantly differed from random. This result can be used to determine the faunal similarity of habitats as well as species habitat preferences.

### Visualisation and grouping of mutually associated species

It is not uncommon for a single species to be associated with multiple others. It is convenient to visualise such cases using an undirected graph, in which the nodes represent the species from the list of associated ones and the edges represent the identified associations (Fig. 3). The size of the node marker will be proportional to the occurrence or the total abundance (or biomass) of the species, and the edge thickness represents the degree of association. This degree can be defined by various methods, including the use of one of the previously described normalised matrix indices or the probability of obtaining the observed joint co-occurrence, which is calculated using the selected distribution probability mass function. If desired, pairs of incompatible species can be added to the graph, indicating positive or negative associations with different edge colors.

**Figure 3.**
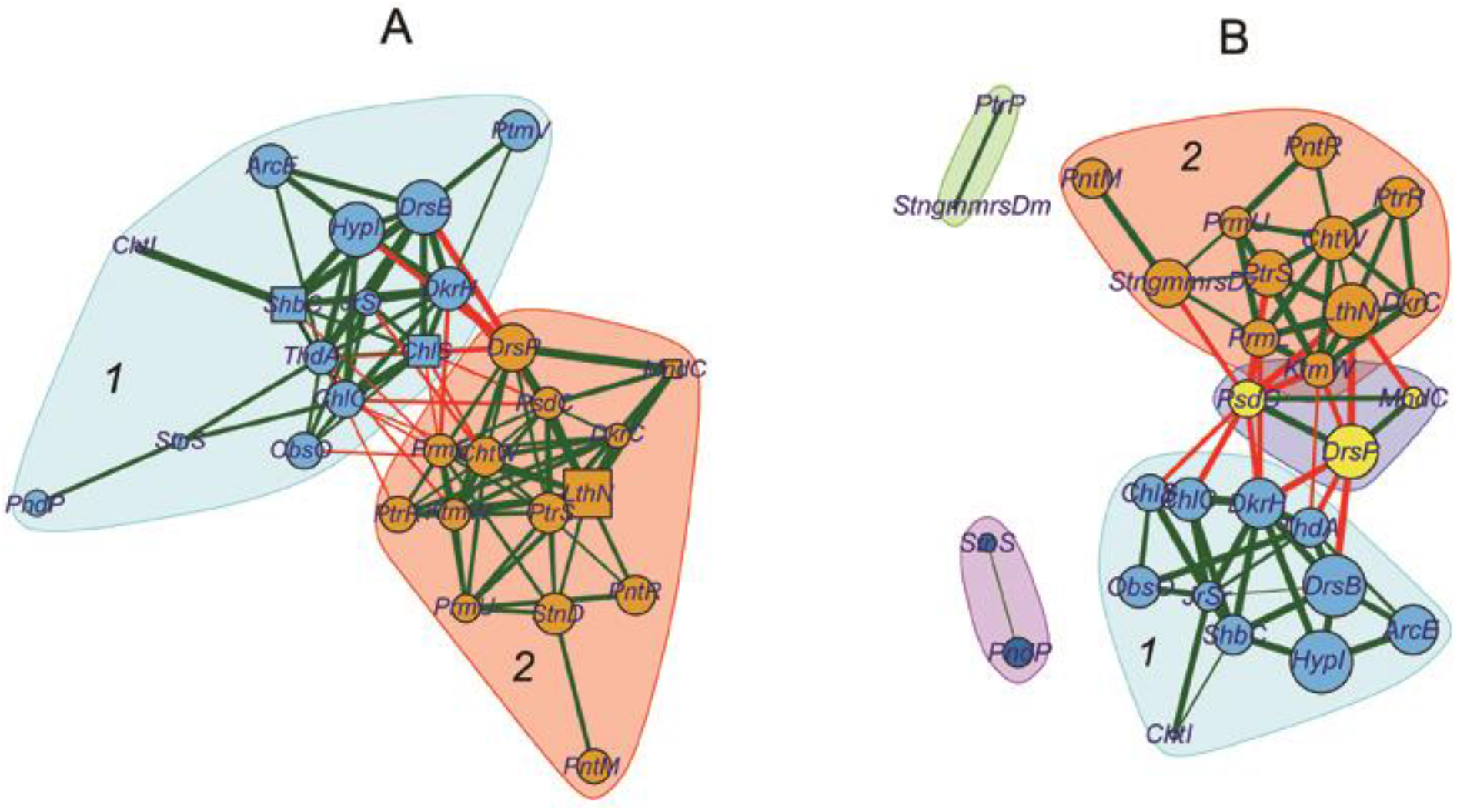
Graph visualisation of positively associated species: A – FE permutation model of the C-score index; B – hypergeometric distribution. The significance level was 0.01. For species abbreviations see (Kurina & Seleznev 2019).

**Figure 4.**
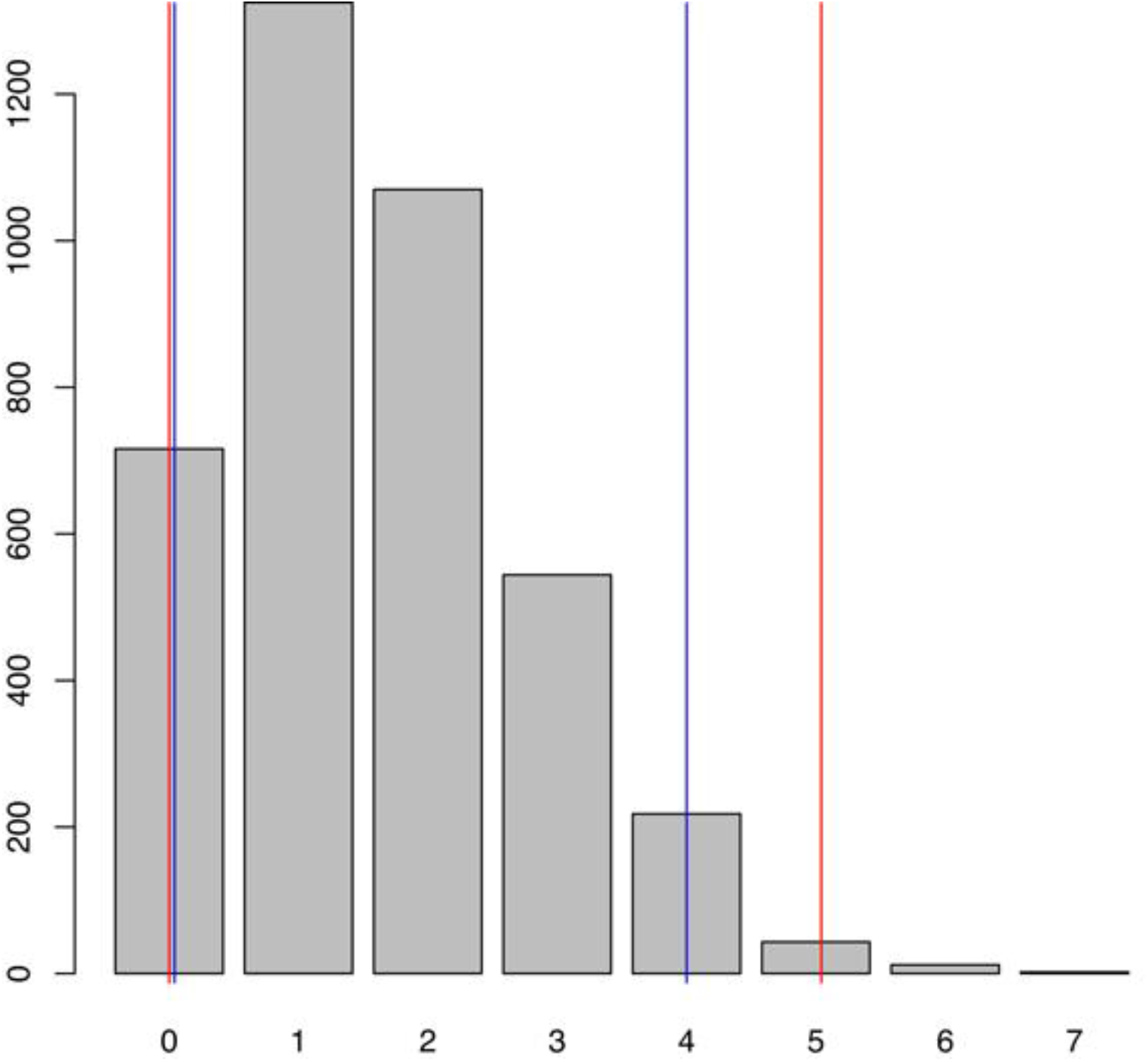
The posterior distribution of the joint co-occurrence of the two species. The blue lines show the limits of the double-sided 90% confidence intervals, and the red lines show the limits of the double-sided 98% confidence intervals.

The layout of graph nodes can be defined using a variety of algorithms, among which we can highlight the Kamada–Kawai algorithm (Kamada & Kawai, 1989), which is based on the physical model of springs (Fig. 2A), and the force-directed Fruchterman–Reingold algorithm (Fruchterman & Reingold, 1991). A further approach is possible, whereby additional meaning is added to the layout of the graph nodes. In particular, one may employ one of the ordination methods based on the frequency of occurrence, total abundance, or biomass of the associated species. In this case, the layout of the nodes will be defined by species scores resulting from the ordination procedure (Fig. 2B).

It is convenient to combine groups of mutually associated species into clusters, which in graph theory terms are referred to as “communities”. The majority of graph node clustering algorithms are based on modularity maximisation, which estimates the extent to which the density of intragroup edges in a given partition is greater than that in a random partition of the same number of edges. There are numerous algorithms of this kind, and they do not require the number of clusters to be prespecified, unlike k-means clustering. In each cluster, the species with the largest number of the strongest connections can be identified as the “core” of the species assemblage, which determines its properties. The species clustering results can be used as input data for the analysis of species assemblages and interactions between species within them.

### Analysis of mutually associated species groups

It is obvious that the selected groups will differ in both the density of connections within them and in the average connection strength. The density of connections (or cluster connectivity) can be calculated as the quotient of the number of realised connections and the number of all possible connections in a cluster: *Cn = 2k/(n·(n−1))*, where *k* is the number of realised connections and *n* is the number of species in the cluster. The average connection strength in the cluster can be estimated as the average of any edge thickness values proposed above or as the average normalised joint co-occurrence *Cf = 2·x/(n+m)*, where *n* and *m* are the number of samples containing the first and second species, respectively, and *x* is the number of samples containing both species together.

Let us reduce the value of the cumulative distribution function *P*_*x*_ to fractions of one. For negatively associated pairs, the value is given by *1− P*_*x*_*/p-value*. For positively associated pairs, the value is given by *1−(1−P*_*x*_*)/p-value*. The total connectivity strength, which characterises both the density and connection strength in the cluster, is calculated as the average value of the distribution function in fractions of one of all significant connections, including zeros for insignificant connections. Consider a cluster of three species in which two pairs of positively associated species have been identified. The significance level for these associations was set at 0.05. The inverse values of the cumulative distribution function of these associations *(1−P*_*x*_) were 0.04 and 0.01, respectively. The total connectivity strength of this cluster is therefore (0.2 + 0.8 + 0)/3 = 0.33. This characteristic is normalised in the range [0,1], thus enabling comparisons between different clusters. Furthermore, it is balanced, in that if there are few statistically significant associations in the cluster, a large number of zeros will affect the total connectivity strength, and conversely, if there are many significant associations, the connections’ strength will affect it more.

### Practical application

The proposed statistical approach was employed for the repeated analysis of data on the occurrence of 37 alien macrozoobenthic species in 255 samples from Volga River reservoirs. Previously, these data were analysed using the C-score index, 1000 FE permutations and a double-sided 98% confidence interval (Kurina & Seleznev, 2019). The results of the comparison of the models are presented in Table 1.

**Table 1.** Comparison of models of associated species: A – C-score index model, B – hypergeometric distribution model with confidence intervals, C – hypergeometric distribution model with Bayesian credible intervals, D – binomial distribution model with confidence intervals, E – binomial distribution model with Bayesian credible intervals. A double-sided 98% confidence interval was used; * – averaged values for 5 simulations.

| Characteristics | Models |  |  |  |  |
| --- | --- | --- | --- | --- | --- |
|  | A | B | C* | D | E* |
| Associated pairs | 78 | 75 | 80 | 53 | 61 |
| Incompatible pairs | 5 | 6 | 5 | 4 | 3 |
| Analysis duration, sec. | 174 | 13 | 30 | 13 | 30 |

Models based on the C-score index and hypergeometric distribution yielded comparable results, although there was a slight discrepancy in the set of species pairs. However, there was a significant difference in the computational resources required for each method. The confidence interval method produces a slightly more conservative result than Bayesian inference in both statistical distribution models. The result of the binomial distribution model is more conservative due to more “heavy tails” in the region of low and high values of *x* (Fig. 1).

Based on the results of the analysis using the permutation model of the C-score index and the confidence intervals of the hypergeometric distribution, we constructed graphs (Fig. 3) and compared them. The thickness of the edges was calculated using the value of the cumulative distribution function *P*_*x*_ reduced to fractions of one, as described above. The node marker size was proportional to the species occurrence frequency. The node layout is generated using the Kamada–Kawai algorithm, while clustering is performed by the multi-level modularity optimisation algorithm (Blondel et al., 2008).

A comparison of the graphs of associated species obtained using the C-score index (Fig. 3A) and the hypergeometric distribution (Fig. 3B) revealed that the latter had greater modularity (0.41 vs. 0.32). This is due to the isolation of some species into separate clusters, while the number of intergroup connections is approximately equal (16 vs. 18). At the same time, two main groups of species remain: deep-water (I) and shore (II) species assemblages. Additionally, three species are separated into an intermediate, depth-independent assemblage associated with sandy sediments. Thus, the new grouping of associated species allows us to identify an additional abiotic factor that affects the spatial distribution of species. The two groups obtained by the two different methods are analysed below (Table 2).

**Table 2.** Characteristics of deep-water (I) and shore (II) clusters of associated macrozoobenthic species selected using the C-score index (C-score) and hypergeometric distribution function (HyperH).

| Characteristics | Methods |  |  |  |
| --- | --- | --- | --- | --- |
|  | C-score |  | HyperH |  |
| Cluster | I | II | I | II |
| Species number, *n* | 14 | 14 | 11 | 11 |
| Associations number, $k$ | 39 | 30 | 27 | 27 |
| Connectivity | 0.36 | 0.47 | 0.49 | 0.49 |
| Average connection strength | 0.92 | 0.95 | 0.86 | 0.77 |
| Total connectivity strength | 0.39 | 0.31 | 0.42 | 0.38 |

The characteristics of the clusters demonstrated that when a hypergeometric distribution was used, there were three pairs of species fewer in both groups, which resulted in an increase in connectivity. Despite a slight decrease in the average connection strength, the total connectivity strength increased.

The aforementioned pair of species, *Katamysis warpachowskyi* (observed in 16 samples) and *Pterocuma rostrata* (observed in 25 samples) occurred together in 5 out of 255 samples. The pair was analysed using different methods. The C-score index for this pair is 220 and does not always fall outside the upper 99% confidence interval of the simulated index distribution. Consequently, the mutual association of these species is on the border of statistical significance. The value of the hypergeometric distribution function, i.e., the probability of obtaining the actual (5) or greater co-occurrence of species for the given parameters (individual occurrence), is 0.0125. This value does not fall outside the upper 99% limit of the confidence interval.

In order to solve the same task, we use Bayesian inference. As a prior distribution, we simulate 10^6^ values for the total number of samples *N*, which are uniformly distributed over the range [25, 255]. We will use the probability mass function of the hypergeometric distribution as a generative function, with the observed individual occurrence of species (25, 16) and the simulated total number of samples. Subsequently, the posterior distribution, which was filtered by the actual number of samples (255), will yielded the distribution of the number of joint co-occurrences of the two species (Fig. 4).

Fig. 4 demonstrates that the observed value (5) does not exceed the upper 99% limit of the confidence interval. Consequently, the application of Bayesian inference with a 2% probability of type I error does not allow us to conclude that the joint co-occurrence of *Katamysis warpachowskyi* and *Pterocuma rostrata* is statistically different from random.

The publication of Veech (2013) represents the results of applying that author’s probabilistic model to 10 different datasets with a significance level of 0.05. Comparisons were made between the published results and those obtained by the methods proposed above on data taken from Galapagos Island finches (Sanderson, 2000), Great Basin Desert rodents (Bowers & Brown, 1982), carabid beetles on islands in lakes of Poland (Ulrich & Zalewski, 2006; Gotelli & Ulrich, 2010), as well as bog and forest ants in New England (Gotelli & Ellison, 2002) (Table 3).

**Table 3.** Comparison of different models of associated species pairs. The number of positive and negative associations is given with a slash. * – rounded results of 5 simulation series; ** – from (Veech, 2013).

| Characteristics | Animals |  |  |  |  |
| --- | --- | --- | --- | --- | --- |
|  | Finches | Rodents | Beetles | Bog ants | Forest ants |
| Species number | 13 | 16 | 71 | 24 | 37 |
| Sites number | 17 | 39 | 17 | 22 | 22 |
| Probabilistic model** | 14/1 | 10/6 | 58/10 | 2/0 | 24/12 |
| C-score index with 1000 FE resamples | 17/0 | 15/1 | 70/11 | 9/0 | 30/12 |
| Hypergeometric distribution, | 14/1 | 11/6 | 61/10 | 12/0 | 32/12 |
| Hypergeometric distribution, credible | 16/1* | 11/6* | 64/10* | 10/0* | 31/12* |
| Binomial distribution, confidence | 0/0 | 3/3 | 7/0 | 8/0 | 10/1 |
| Binomial distribution, credible intervals | 0/0* | 3/3* | 6/0* | 8/0* | 11/1* |

The results of the probabilistic model and the hypergeometric distribution model are found to be highly correlated, with the FE resampling null model exhibiting a tendency to overestimate the number of positively associated pairs and underestimate the number of negatively associated pairs. The only exceptions to this are the bog and forest ant datasets, for which both the probabilistic model and the FE resampling null model were found to be more conservative than the hypergeometric distribution model. It is also worth noting that the binomial distribution model is the most conservative of all those considered.

### Ecological applications

In order to ensure the successful using of the proposed methodology in future investigations, it is essential to have a clear understanding of the ecological meaning of species falling into an associated pair or group. To date, the methodology has been employed in six studies, three of which were based on macrozoobenthos data (Kurina et al., 2019; Prokin et al., 2021; Kurina et al., 2022) and three on zooplankton data (Kotov et al., 2022; Chertoprud et al., 2023; Andreeva et al., 2023). To identify associated pairs of species, the initial data was filtered in various ways in all studies except (Prokin et al., 2021), thus allowing us to discuss analyses of taxocoenoses rather than complete biological communities.

In all our studies, the principal criteria for categorising species into associated groups were environmental factors, such as depth, current velocity, salinity and other physico-chemical parameters of the water, the presence of water plants, the type of bottom sediment or the type and level of anthropogenic influence. It is likely that the common requirements of environmental factors also explain the occurrence of ecosystem engineers such as the quagga mussel *Dreissena bugensis* (Andrusov, 1897) and the zebra mussel *D. polymorpha* (Pallas, 1771) with connected species in the associated groups (Kurina et al., 2019; Kurina et al., 2022).

A preliminary analysis of the results obtained thus far suggests that the observed associations between species indicate a similar selectivity to a set of environmental factors that limit their distribution at the scale of a particular study. However, changes in the area or scale of the study may result in considerable alterations to the hierarchy and importance of factors and consequently, the groups of associated species. To test and refine this assumption, further studies on data from different groups of organisms, as well as meta-analyses of these studies, are required.

### Sofware

The senior author’s open access web application (http://apps.ibiw.ru/coobs) implements the described methodology and allows users to obtain results in text and graphic formats. As an analysis method, hypergeometric or binomial distributions can be selected. The choice of pairs of associated species is performed using confidence intervals or Bayesian credible intervals of selected distribution at the given significance level. In the case of Bayesian credible intervals, 10^6^ uniformly distributed total number of samples is used as a prior distribution, with the individual species occurrences fixed. An undirected graph is constructed by several methods, using positive or negative associations only or using all selected associations. To group graph nodes, various algorithms are used that take into account the strength of the connections between graph nodes as a positive edge weight factor. In addition, it is possible to highlight the “Spin glass” algorithm (Traag & Bruggeman, 2008), which is capable of handling negative associations. This algorithm is able to work correctly with negative edge weights, in contrast to other methods that consider them to be null. The “Optimal community structure” algorithm (Brandes et al., 2008) is also worthy of note because it solves the graph node grouping task as an integer programming problem and builds a graph with maximum modularity. However, this method also fails to consider negative associations, such as species incompatibility, and has exponential complexity. This is because the task of maximising modularity appears to belong to the NP class (Brandes et al., 2006).

The application was developed in the R 4.2 statistical analysis environment using the package igraph (Csardi & Nepusz, 2006). The web interface was designed using the Shiny package (Chang et al., 2021).

## Conclusion

The proposed methodology differs from those that use permutation null models in its simplicity of implementation and in its low requirements for computing resources. This approach allows to obtain reproducible and easily interpreted results with a given significance level in the form of a list of pairs of mutually associated or incompatible species. This list can later be visualised and analysed both mathematically and in terms of studying the ecological characteristics of individual species and the features of the formation of species assemblages and communities.

One disadvantage of the methodology is that it uses only qualitative data on the presence or absence of species in the samples, without considering their quantitative characteristics, such as total abundance or biomass. It is possible that two species may occur in samples together rarely but in large numbers and separately often but in small numbers (or vice versa). However, this phenomenon is not detected when discrete distributions are used. To account for quantitative data, it seems that the use of continuous counterparts of discrete distributions or some other methods will be required.

## Acknowledgements

The authors are grateful to D.Y. Medvedeva and P.N. Petrov (Moscow) for their help with improving the English of the manuscript and V.K. Shitikov (Tolyatti) for a constructive discussion of this study.

